# Prophages are associated with extensive CRISPR-Cas auto-immunity

**DOI:** 10.1101/2020.03.02.973784

**Authors:** Franklin L. Nobrega, Hielke Walinga, Bas E. Dutilh, Stan J.J. Brouns

## Abstract

CRISPR-Cas systems require discriminating self from non-self DNA during adaptation and interference. Yet, multiple cases have been reported of bacteria containing self-targeting spacers (STS), i.e. CRISPR spacers targeting protospacers on the same genome. STS has been suggested to reflect potential auto-immunity as an unwanted side effect of CRISPR-Cas defense, or a regulatory mechanism for gene expression. Here we investigated the incidence, distribution, and evasion of STS in over 100,000 bacterial genomes. We found STS in all CRISPR-Cas types and in one fifth of all CRISPR-carrying bacteria. Notably, up to 40% of I-B and I-F CRISPR-Cas systems contained STS. We observed that STS-containing genomes almost always carry a prophage and that STS map to prophage regions in more than half of the cases. Despite carrying STS, genetic deterioration of CRISPR-Cas systems appears to be rare, suggesting a level of escape from the potentially deleterious effects of STS by other mechanisms such as anti-CRISPR proteins and CRISPR target mutations. We propose a scenario where it is common to acquire an STS against a prophage, and this may trigger more extensive STS buildup by primed spacer acquisition in type I systems, without detrimental autoimmunity effects. The mechanisms of auto-immunity evasion create tolerance to STS-targeted prophages, and contribute both to viral dissemination and bacterial diversification.

## INTRODUCTION

Clustered regularly interspaced short palindromic repeats (CRISPR) and CRISPR-associated proteins (Cas) are defense systems, which provide bacteria and archaea with an adaptive and heritable immunity against invading genetic elements such as bacteriophages or plasmids (1–3). Immunity is conferred by small sequences, known as spacers, which are taken up from the invaders’ genome and integrated into the CRISPR locus (2). At the CRISPR locus, spacers function as the system’s memory, and are used in the form of guide RNA to specifically recognize and degrade foreign DNA or RNA (3–5). While known to be highly specific for their target, CRISPR-Cas systems do pose a risk for auto-immunity if spacers from the host chromosome are mistakenly acquired (6). These self-targeting spacers (STS) have been reported in numerous species, and their most likely consequence is cell death by directing cleavage and subsequent degradation of the host genome (7,8). Escape from the lethal outcome of auto-immunity occurs for cells selected for mutations on the target sequence (9,10) and/or for inactivation of CRISPR-Cas functionality via, for example, mutation or deletion of the Cas genes, spacers, repeats, or protospacer adjacent motifs (PAM). The action of anti-CRISPR (Acr) proteins encoded by prophages may also prevent auto-immunity (11). In fact, the presence of STS in a genome has been suggested (11,12) and recently successfully employed (13) as a strategy to discover new Acrs.

Auto-immunity has been mostly regarded as a collateral effect of CRISPR-Cas systems, but it has also been suggested to play a role in the evolution of bacterial genomes on a population level by influencing genome remodeling (9). Although reported only on isolated examples, CRISPR-Cas systems have been speculated to act like a regulatory mechanism (14–17). Auto-immunity has also been proposed to be triggered by foreign DNA with similarity to the bacterial chromosome (18).

Here we take a closer look at STS in the many types and subtypes of CRISPR-Cas systems to identify the incidence, distribution and mechanism of evasion of potential CRISPR-Cas auto-immunity in bacteria. We demonstrate that STS are frequently observed in bacterial genomes, and that bacteria have evolved mechanisms to evade death by auto-immunity while preserving their CRISPR-Cas systems. We propose that the integration of phages in the bacterial chromosome provides evolutionary advantages to the bacteria (e.g. acquisition of virulence traits) but is also the primary trigger of STS acquisition in CRISPR arrays. We further suggest that mechanisms of evasion from auto-immunity create tolerance to the integrated invaders, benefiting both bacteria and phage populations by allowing the acquisition of novel genetic information by the bacteria, and by promoting phage (passive) dissemination in the bacterial population.

## Material and methods

### Detection of CRISPR arrays

The complete genome collection of the PATRIC database (19) (a total of 110,334 genomes) was used in our analysis. CRISPR arrays were predicted for each genome using CRISPRDetect 2.2.1 (20) with a quality score cut-off of 3.

### Detection of self-targeting spacers

All spacers were blasted (blastn-short option, DUST disabled, e-value cut-off of 1, gap open, and gap extend penalty of 10) against the source genome. The blastn results were filtered for a minimum identity higher than 90% with the target. Any hit on the genome was considered a self-target, except for those within all of the predicted CRISPR arrays, including arrays identified with a CRISPRDetect quality score below 3. Hits closer than 500 bp from each end of the predicted arrays were also ignored to avoid considering spacers from the array that were possibly not identified by CRISPRDetect. Spacers with flanking repeats of identity score lower than 75% to each other were discarded as they may have been erroneously identified as spacers. Of these, only spacers smaller than 70 bp and a repeat size between 24 and 50 bp were retained in the dataset. Finally, STS from CRISPR arrays of two or fewer spacers were excluded, except when the associated repeat belonged to a known CRISPR repeat family, as identified by CRISPRDetect. Duplicates were removed by search of similar genomes, contigs and arrays.

### Classification of CRISPR-Cas systems

The CRISPR-Cas systems of STS-containing genomes were classified using MacsyFinder (21) in combination with Prodigal (22), and the CRISPR-type definitions and Hidden Markov Models (HMM) profiles of CRISPRCasFinder (23). The classification of the repeat family of the CRISPR array was obtained using CRISPRDetect. Genomes carrying two or more CRISPR-Cas types were labeled as ‘mixed’, and those having CRISPR-Cas arrays but no *cas* genes were labeled as ‘no Cas’. Systems which could not be assigned a CRISPR sub-type and which were missing at least one *cas* gene (but contained no less than one *cas* gene) were classified as ‘incomplete’. The final classification of each genome can be found in Supplementary Table S1.

### Analysis of the genomic target

The orientation of the arrays was determined by CRISPRDetect using the default parameters of CRISPRDirection. After this, the STS sequence was used for a gapless blastn at the target and to retrieve the PAM downstream or upstream of the STS based on the CRISPRDetect classification (see Supplementary Table S1). The targets were then analyzed for the correct PAM sequence by comparison with the expected PAM for the different CRISPR-Cas types as previously described (11,24,25). The consensus PAM sequences used in this analysis are shown in Supplementary Table S2. Genes of STS-containing genomes were predicted using Prodigal and annotated using Interproscan (26) and Pfam (27) domain prediction. Prophage regions in the genomes were detected using VirSorter (28), and used to identify STS targeting these regions. Transposons were also detected in the genomes using Interproscan (26) (Supplementary Table S3). Targets of the STS with e-value <10^−5^ were grouped by function to identify possibly enriched hits separately for prophage and endogenous regions. Only those hits associated with predicted correct PAMs were subjected to this analysis.

### Distance between self-targeting spacer and prophages

Contigs predicted to contain prophages were extracted and used to create a hit density map based on STS distance to prophage(s).

### Identification of anti-CRISPR proteins

The amino acid sequences of known Acrs (29) were used for homology search in the STS-containing genomes using BLASTp with an e-value limit of 10^−5^.

### Statistical analysis

A binomial test was performed on CRISPR arrays of different sizes to test the hypothesis that STS at the leader side of the CRISPR array are more common. Only STS from CRISPR arrays smaller than 50 spacers were considered because larger arrays are too scarce to result in a reliable statistical analysis. A chi-squared test was used to determine statistical significance between percentages of populations. Statistical significance was considered for *P* < 0.05.

### Software

GNU parallel was used to parallelize tool runs and for parsing of output files (30). Biopython package (31) functions were used for specific analysis, such as GFF parser for prodigal files, pairwise2 for removing false positives based on repeat identity, and nt_search for matching of the PAM. All data collected was managed using Python package Pandas (32). Python packages SciPy (33), Matplotlib (34) and Seaborn (35) were used for statistical analysis and visualization.

## Results

### Self-targeting spacers (STS) are often found in CRISPR-encoding bacteria

We scanned 43,526 CRISPR-encoding genomes for spacers with >90% sequence identity to the endogenous genomic sequence that is not part of a CRISPR array. We decided upon this definition of STS as a 10% mismatch between spacer and target can still trigger a functional CRISPR response (direct interference and/or priming in type I) in many CRISPR-Cas types (36–41). For clarity, we note that our definition of STS may exclude or include certain sequences as a result. For example, STS protospacers that suffered extensive mutations may be excluded, while spacers that target non-genomic regions of high similarity to a genomic region may be included. We found that 23,626 out of 1,481,476 spacers (1.6%) are self-targeting based on this cutoff. Approximately half of those (12,121, 0.8%) had 100% sequence identity to the genome from which the spacers were derived (frequency of STS with mismatches can be seen in Supplementary Table S4), a percentage higher than previously reported (0.4% with 100% identity) (14). Similar to previous observations with smaller datasets (14), about one fifth (19%, 8,466) of CRISPR-encoding genomes have at least one STS in one of their CRISPR arrays.

We further looked into how frequent STS were in different types of CRISPR-Cas systems (Figure 1A). STS were detected in genomes containing CRISPR-Cas systems of almost all subtypes, and were more prevalent (>40%) in CRISPR-Cas types I-B and I-F. Curiously, genomes containing STS are almost absent in type III-A, but present between 10 and 20% in type III-B, C and D systems. Moreover, length of the STS agreed with reported preferred spacer length for different CRISPR-Cas subtypes (Supplementary Figure S1) (42–44).

**Figure 1.**
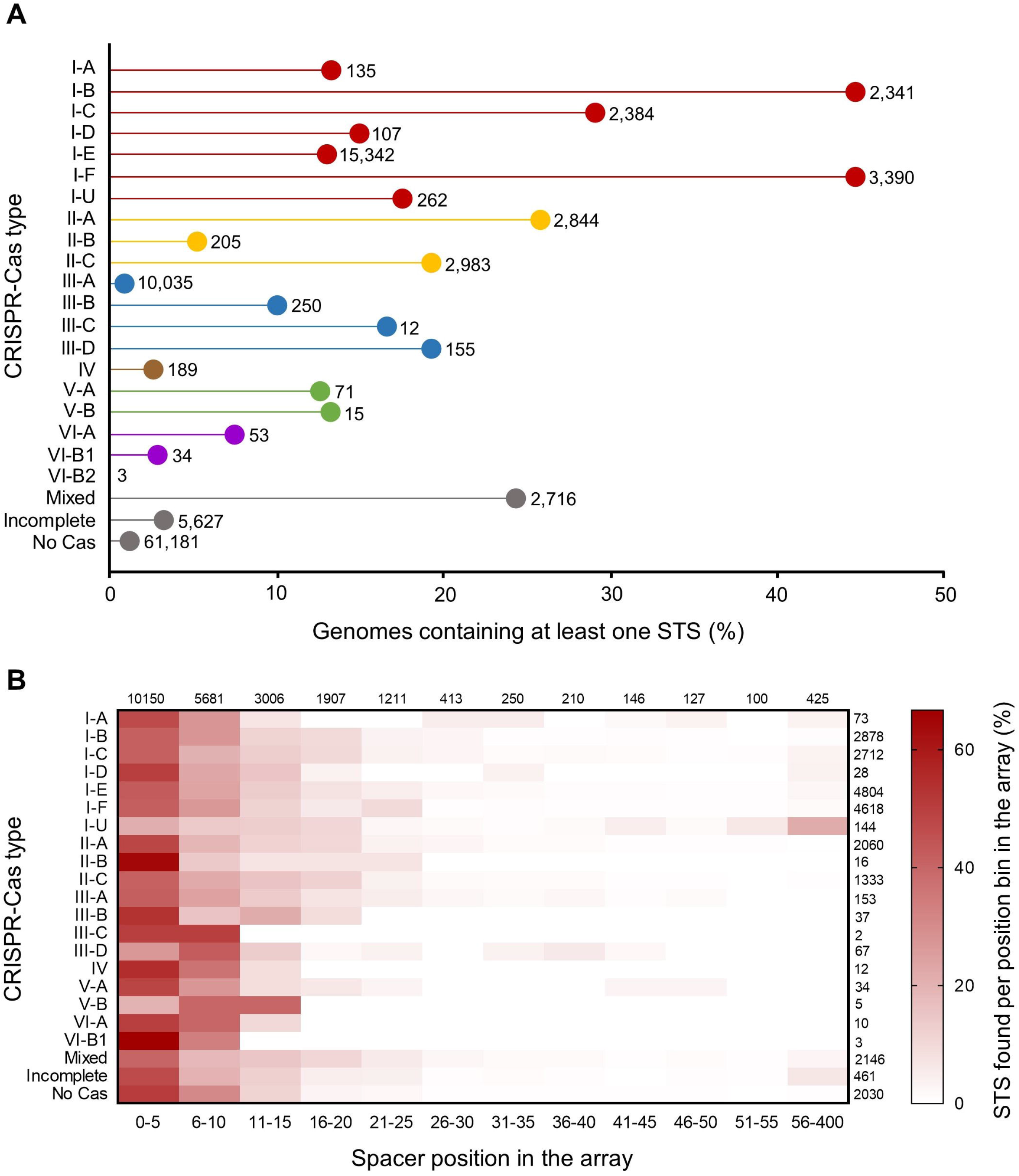
Self-targeting spacers (STS) in CRISPR-containing bacteria. (**A**) Frequency of genomes containing STS for the different subtypes of CRISPR-Cas systems. Total number of CRISPR-containing genomes analyzed is given for each row; (**B**) Heatmap of STS position in the CRISPR array for each CRISPR-Cas subtype, using corrected orientation of the CRISPR arrays. Scale bar represents percentage of STS found per position bin in the CRISPR array. Total number of STS analyzed per CRISPR-Cas subtype is given for each row, while total number of STS per position bin is given for each column.

It has been suggested that following the integration of an STS, the CRISPR-Cas system must become inactivated in order to survive, and that this phenomenon could explain the abundance of highly degraded CRISPR systems that contain *cas* pseudogenes (14). Recent experimental evolution studies have shown that large genomic deletions encompassing the entire CRISPR-Cas locus can occur as a consequence of auto-immunity to prophages (45). We observed that 12% (979 of 8,466) of the STS-containing genomes contain incomplete CRISPR systems or no *cas* genes, while 88% (7,490 of 8,466) seem to carry intact CRISPR-Cas systems (*P* < 0.0001, chi-squared t-test, Figure 1A). This suggests that CRISPR-Cas deletion can occur as a mechanism to survive STS, but self-targeting can also be overcome through other mechanisms. To note that our homology-based analysis cannot account for small inactivating mutations in *cas* genes that could also render a CRISPR-Cas system non-functional, but we expect that the effect of such recent pseudogenization is minor as inactive pseudogenes tend to be rapidly lost from the genome (46,47). Moreover, we found that most STS locate in the leader proximal positions of the array (Figure 1B, Supplementary Figure S2), with several STS also found in middle and leader distal positions (Figure 1B). To account for potential bias introduced in this analysis by smaller arrays, we generated the same plot for arrays of 10 or less spacers (Supplementary Figure S3). The same trend is apparent, confirming that STS preferably locate near the leader but are also present in later positions in the array. This suggests that the CRISPR system (or at least memory acquisition) remains active after integration of an STS into the CRISPR array and the cell remains viable. Correct CRISPR array orientation prediction remains challenging in some cases (48), and there may be some arrays in our database whose orientation was predicted incorrectly by the CRISPRDirection tool. This may lead to noise in the positionality of STS. Still, we are confident on our overall observations as CRISPRDirection is backed up by experimental evidence for most CRISPR types, including type I-U (49).

In summary, STS are common among bacteria harboring all types of CRISPR-Cas systems, but especially types I-B and I-F. Importantly, STS-containing bacteria seem to preserve CRISPR-Cas, perhaps by employing alternative mechanisms to avoid the lethal effects of auto-immunity.

### STS are enriched in prophage-containing genomes

To understand if targeting of endogenous regions by STS could have a regulatory role in gene expression, we looked at the position of STS hits in the genome and determined if these were in coding or non-coding regions. In general, no preference for targeting non-coding regions was observed, with coding regions being predominant in most types of CRISPR-Cas systems (*P* < 0.05, chi-squared test, Supplementary Table S5), with the exception of STS of types I-D, III-A, III-B, V-B and VI-A CRISPR-Cas systems for which intergenic and coding regions are equally targeted (*P* > 0.05, Figure 2A, Supplementary Table S5). This suggests that there is no apparent link between CRISPR-Cas auto-immunity and regulating promoter activity for gene expression. Still, no absolute conclusions can be drawn about a potential regulatory role of STS since direct targeting of genes (coding regions) leads to programmed regulation of gene expression (50–53). Also, in most cases we could not detect a preference for targets on the sense or antisense DNA strands (*P* > 0.05, Figure 2A, Supplementary Table S5).

**Figure 2.**
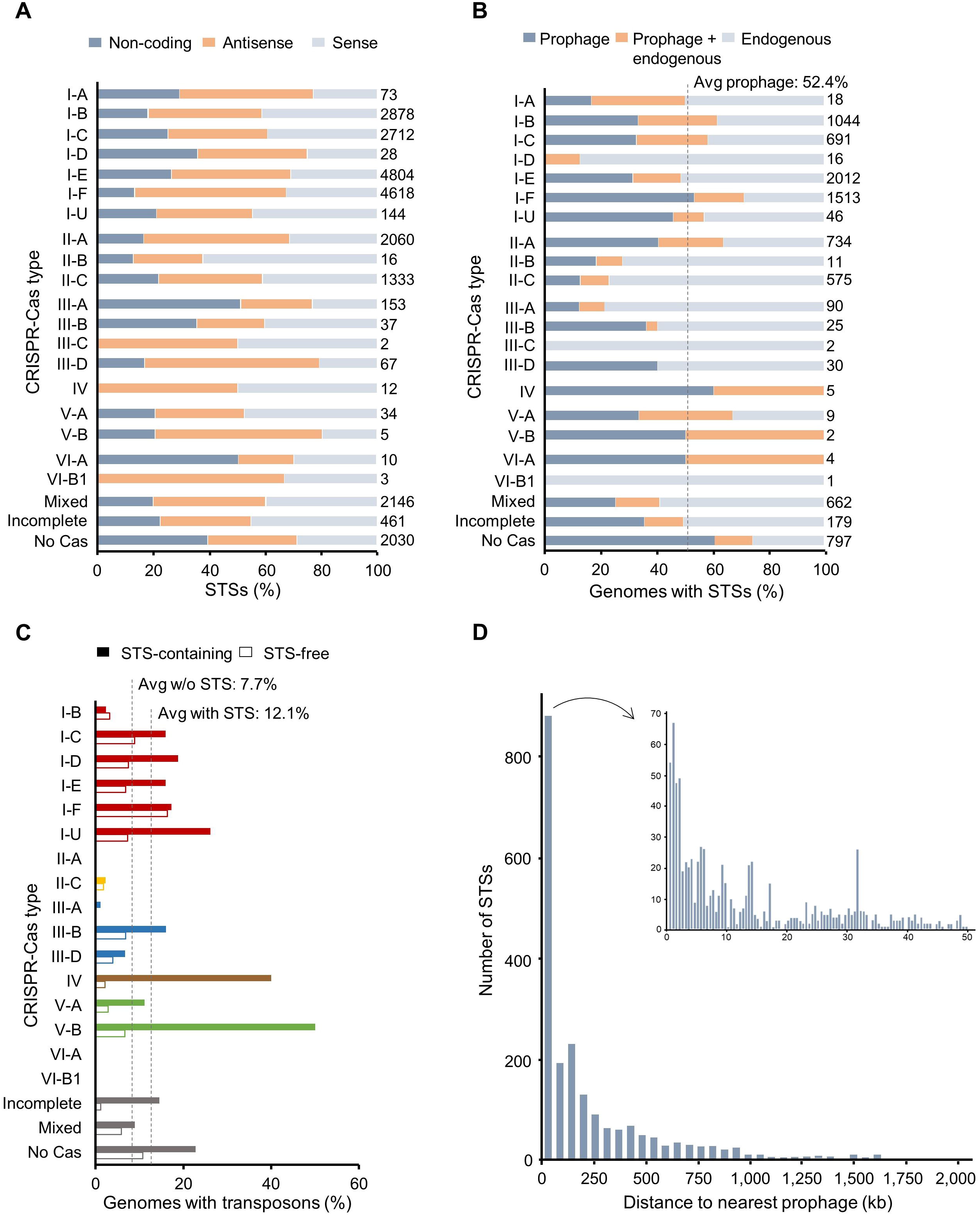
Genomic targets of self-targeting spacers (STS). (**A**) Preference of STS for targeting sense or antisense strands of coding regions, or non-coding regions of the bacterial genome. Values were normalized to the percentage of coding or non-coding regions of the genome. Total number of STS are indicated at the end of bars; (**B**) Prevalence of STS targeting only prophage regions, endogenous genomic regions, or both, in each CRISPR-Cas subtype. Total number of STS-containing genomes are indicated for bars; (**C**) Transposon abundance in STS-containing genomes (full bars) and STS-free genomes (empty bars) for each CRISPR-Cas subtype; (**D**) Distribution of distances between STS protospacer and the nearest prophage. Internal plot shows the largest peak binned into smaller (0.5 kb) increments.

**Figure 3.**
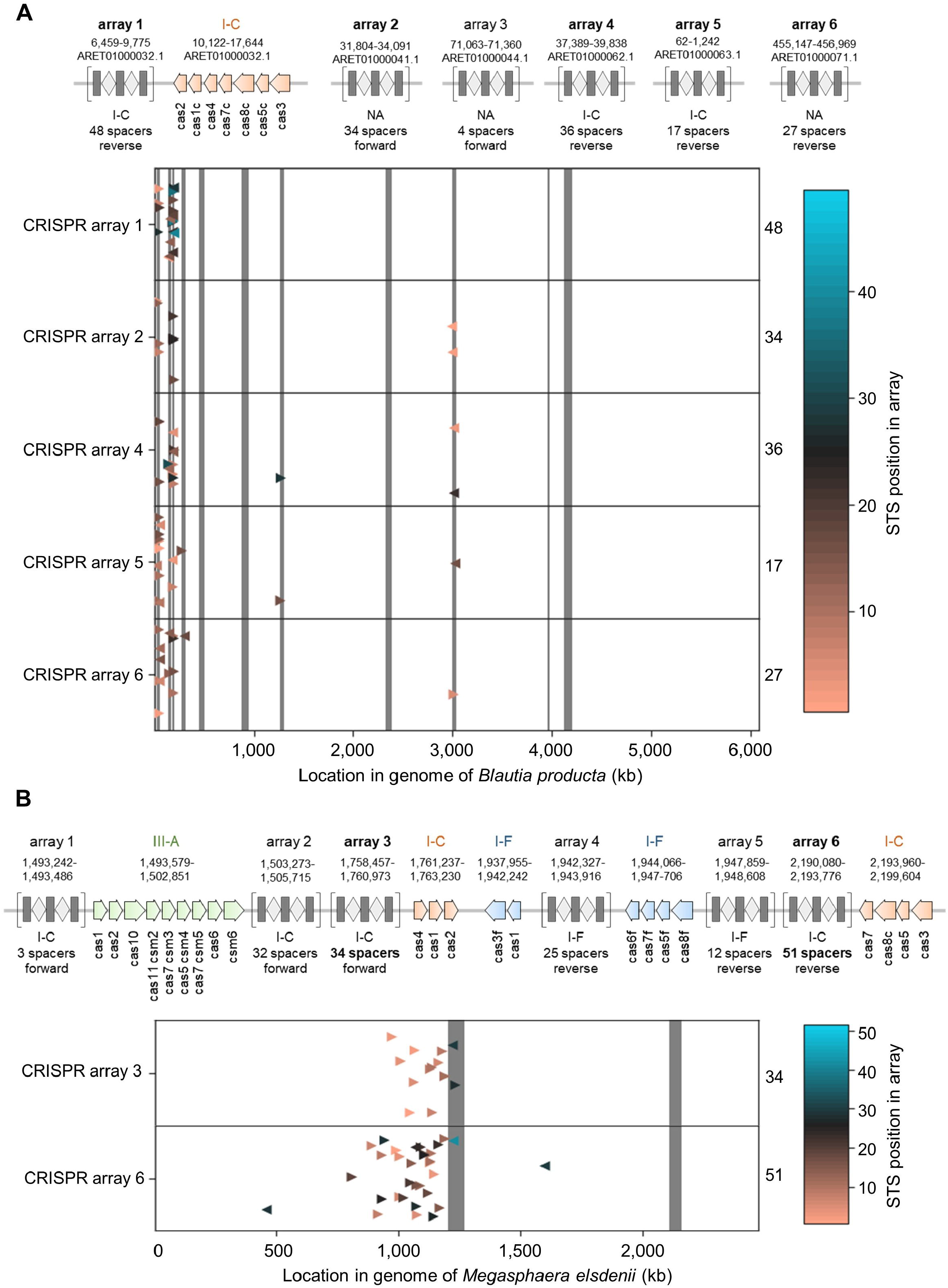
Extreme cases of self-targeting in prophage regions of bacterial genomes containing a high number of STS with 100% sequence identity to the target. (**A**) *Blautia producta* strain ATCC 27340 (accession number ARET01000032) carries a type I-C CRISPR-Cas system and 11 prophages, and has 162 STS. Arrays identified in different contigs from where STS originate are represented in the y-axis; and (**B**) *Megasphaera elsdenii* strain DSM 20460 (accession number NC_015873) carries types I-C, I-F and III-A CRISPR-Cas systems and two prophages, and has 85 STS. STS originate from two out of six CRISPR arrays (array 3 at 1,758,457-1,760,973 bp, and array 6 at 2,190,080-2,193,776 bp), which are associated with the type I-C system and are represented in the y-axis. For both panels, prophage regions are denoted in dark grey, STS hits are represented as colored triangles, and scale represents position of STS in the array. The total number of STS per contig or array is shown for each row.

Bacteriophages are common targets of CRISPR-Cas systems and exist abundantly in nature. Because some bacteriophages can integrate into the bacterial chromosome, we next investigated if the presence of prophages in a genome would associate with the presence of STS. We identified prophage regions in the STS-containing genomes and observed that, on average, 52.4% of the STS-containing genomes have STS with protospacers in prophage regions, with type I-F CRISPR-Cas systems showing up to 70% genomes with prophage hits (Figure 2B). Interestingly, we also observed that 96.9% (8,203 out of 8,466) of the STS-containing genomes have at least one integrated prophage, while only 28.5% (9,992 out of 35,060) of the STS-free genomes contain prophages (*P* < 0.0001, chi-squared test). It therefore appears that STS is linked to carrying prophages.

We further questioned if STS were also enriched in bacteria containing other mobile genetic elements able to integrate into the bacterial genome. To do so, we looked at the prevalence of transposons in STS-containing and STS-free genomes of bacteria with CRISPR arrays. We observed a moderately higher prevalence of transposons in STS-containing genomes (12.1% vs 7.7%, or 10.9% vs 5.0% when discarding incomplete and no Cas genomes, *P* = 0.004 and *P* = 0.001, respectively, chi-squared test) (Figure 2C).

We next wondered if collateral targeting of prophage regions would lead to STS of endogenous genomic regions flanking the prophage. To test this we mapped the distance of STS in the genome to the nearest prophage region. For this we considered only STS targeting regions of complete genomes and contigs which contained a prophage. 59.5% of these STS target a prophage region, while the remainder mostly target the nearby endogenous genome (Figure 2D). Distances to prophage were also normalized by contig length to discard possible variations due to differences in contig size, which shows a similar pattern of STS hitting regions close to the prophage (Supplementary Figure S4). This suggests that targeting of endogenous regions is indeed related to proximity to a prophage region. As the definition of prophage boundaries may be associated with a certain level of inaccuracy, nearby STS protospacers may also be part of the prophage itself. Because genomic regions flanking prophages are often excised together with the prophage, it is also possible that such regions are subjected to spacer acquisition when the prophage enters its lytic cycle. Finally, prophages tend to repeatedly integrate in the same regions of bacterial genomes, so it is possible that proximal prophage regions are enriched in degenerated prophages as well. All these processes could contribute to the enrichment of STS in prophages and their proximal genomic regions, as shown by our results.

In summary, 63% of STS are linked to prophages or the nearby endogenous genome (<50 kb, see Figure 2D). Thus, our data suggest that the occurrence of STS is strongly linked to the presence of prophages in the bacterial chromosome.

### Interference-functional STS with consensus PAM are frequent in type I CRISPR-Cas systems

To explain how STS are tolerated we first looked at the targeting requirements of CRISPR-Cas systems. In many CRISPR-Cas systems, the correct identification of the target is dependent on a small 2-6 base pair motif immediately adjacent to the target DNA sequence, known as the PAM (54). The PAM is essential for binding to and cleavage of the target DNA by the Cas nucleases, and mutations in this sequence can abrogate targeting (55). To understand how often STS protospacers have a consensus PAM, and can therefore be efficiently targeted, we compared the PAM sequence of the STS protospacer with the expected PAM sequence for the different CRISPR-Cas types previously described (Supplementary Table S2) (24,25,56). We observed that 22.4% of all STS (4,140 of 18,483 STS with 90% sequence identity) and 23.9% of STS with 100 % identity (2,294 of 9,605) have a consensus PAM (Figure 4A and Supplementary Table S6), suggesting these to be functional for direct interference. Type I CRISPR-Cas systems, especially types I-B (29.5%), I-C (44.7%) and I-E (37.0%) have more STS with a consensus PAM (average 27.5%) than type II (average 0.1%) or type V (average 12.8%) (Figure 4A). This may suggest that bacteria encoding type II and type V systems avoid the lethal effects of auto-immunity by having non-functional STS, while bacteria encoding type I systems may employ other evasion mechanisms to withstand the lethal auto-immunity effects of interference-functional STS.

Several factors should be considered when analyzing the role of PAM sequences in tolerance mechanisms to STS. First, the full diversity of functional PAM sequences in nature currently remains unknown, as does their distribution across taxa. Second, PAM sequences can vary widely even within a CRISPR subtype (e.g. in different species) (54,57-59). Third, different CRISPR class I (type I, III and IV) systems may use different PAM sequences for spacer acquisition and for targeting (60). Our analysis has revealed a range of candidate bacteria that can contain mechanisms allowing them to remain viable while carrying interference-functional STS with known consensus PAM sequences. It will be interesting to see these mechanisms further unraveled in future studies.

### Acrs are more prevalent in bacteria carrying STS

To understand how bacteria are able to survive STS while keeping their *cas* genes intact, we assessed the presence of Acrs encoded by prophages. By inhibiting the activity of the CRISPR-Cas system using a variety of mechanisms (reviewed in (29)), Acrs can prevent the lethal effects of STS auto-immunity. In fact, STS have been used to identify new Acr proteins (13,61). We mapped Acrs in the STS-containing genomes using homology searches with all currently known Acrs (29). Acrs were found at low frequency (10.9% average, Figure 4B) but still at levels significantly higher than those found in STS-free, CRISPR-containing genomes (0.3% average, *P* < 0.0001, chi-squared test). The levels of Acrs here reported are a lower bound, as unidentified Acrs may be present in these genomes and these proteins may thus have a higher influence in escaping auto-immunity. Even so, we found many Acr homologs in STS-containing bacteria carrying single type I-B, IV or VI-A CRISPR-Cas systems, for which no Acrs have yet been described (Figure 4B and Supplementary Table S7). Putative Acrs for type I-B and type IV CRISPR-Cas systems were recently identified by using a bioinformatics pipeline (61), but to our knowledge none has yet been suggested for type VI-A.

**Figure 4.**
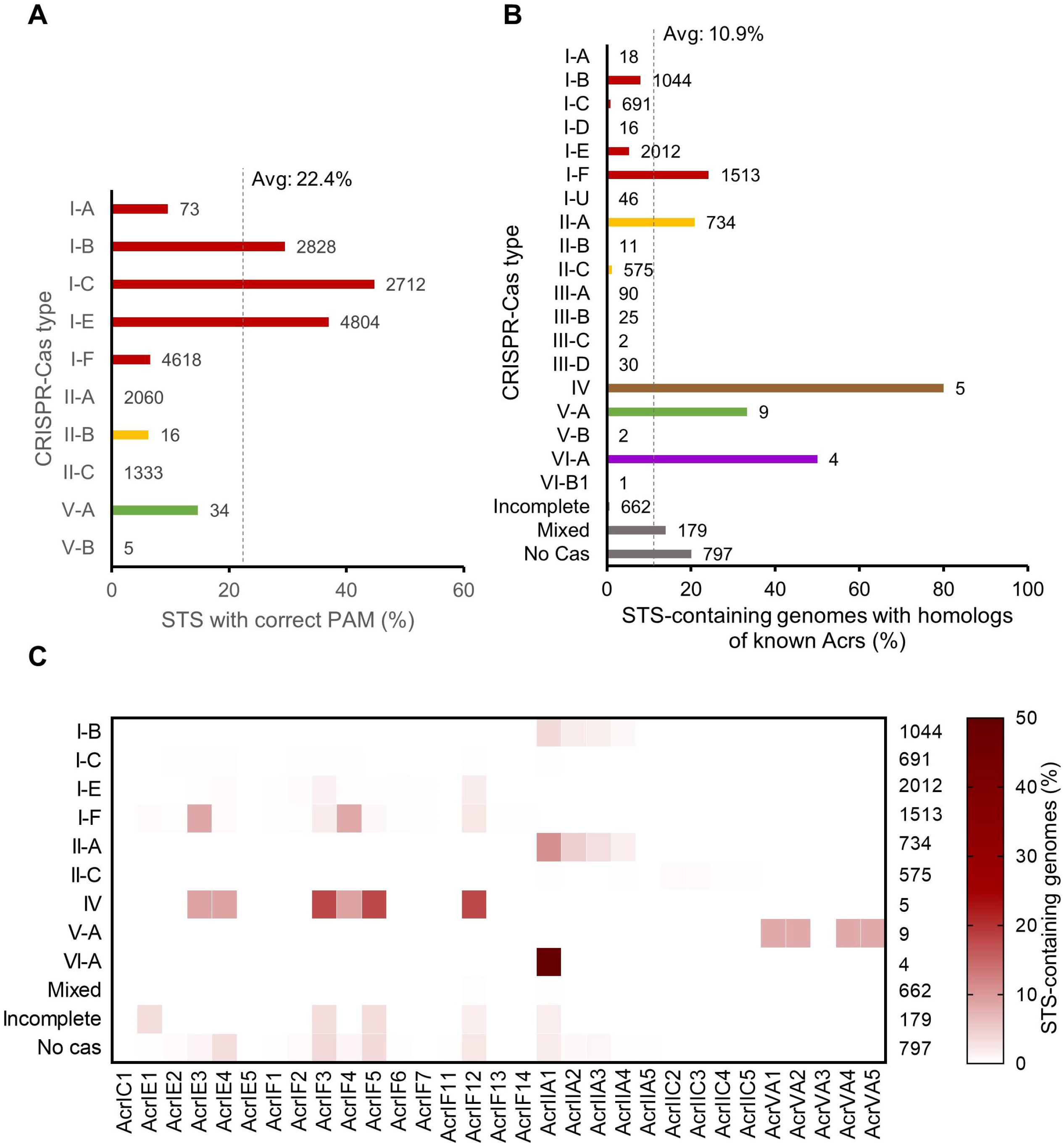
Mechanisms of escape from auto-immunity. (**A**) Levels of self-targeting spacers (STS) associated with correct or incorrect protospacer adjacent motif (PAM) for different types of CRISPR-Cas systems. Only CRISPR-Cas systems with unquestionable type classification and of known PAM were considered. Dashed line indicates the average percentage of STS-containing genomes with correct PAM across CRISPR types; (**B**) Prevalence of STS-containing genomes with Acrs, as found by homology search to known Acrs. Dashed line indicates the average percentage of STS-containing genomes with Acr across CRISPR types; (**C**) Heatmap of prevalence of Acr families in different types of CRISPR-Cas systems in STS-containing genomes. Scale bar represents percentage of STS-containing genomes with a given CRISPR type (row) that contained a homolog of the Acr (column). The total number of STS-containing genomes of each CRISPR-Cas type is given at the end of each row.

Among the newly found Acrs, homologs of AcrIF2-7, AcrIF11-13 and AcrIIA1-4 were the most common in STS-containing genomes (Figure 4C). Interestingly, homologs of AcrIF1-14, AcrIE1-5, and AcrIIA1-4 were found in genomes of diverse CRISPR-Cas subtypes, while homologs of AcrVA1-5 and AcrIIC2-5 appear only in genomes containing the corresponding CRISPR-Cas subtype. Particularly, homologs of AcrIF1-14 and AcrIE1-5 were found in type I and type IV CRISPR-Cas types, while homologs of AcrIIA1-4 were detected in type I, II and VI-A CRISPR-Cas systems. Acr homologs of families that do not correspond to the CRISPR-Cas system found in the bacteria were also recently reported (61). It is possible that some Acr homologs have activity against multiple types of CRISPR-Cas systems, which may occur if the mechanism of inhibition of the Acr is compatible with the multiple types. The ability of Acrs to inhibit different types of CRISPR-Cas systems has already been revealed for some Acrs (62,63), although the specific mechanisms of inhibition have not yet been described.

Anti-CRISPR associated (aca) genes were also found, especially in types I-E and I-F CRISPR-Cas systems, and with higher prevalence of aca1 (488) and aca4 (220) genes (see Supplementary Figure S6 and Supplementary Table S7).

In conclusion, among genomes with a CRISPR system, Acrs are more prevalent in genomes containing STS than in genomes without STS, and it therefore is likely that Acrs play a major role in auto-immunity evasion.

### Amplified self-targeting in prophages regions

In our analysis, we found 1,224 genomes with a number of STS higher than the average (2.5 ± 2.9 STS, Supplementary Figure S5). We decided to take a closer look at two extreme cases and investigate how STS with 100% identity were distributed in the bacterial chromosome (Figure 3). The genome of *Blautia producta* strain ATCC 27340 contains a type I-C CRISPR-Cas system and 11 prophage regions in the chromosome (Figure 3A). This strain contains a stunning 162 STS mostly hitting prophage regions. The genome of *Megasphaera elsdenii* strain DSM 20460 contains three distinct CRISPR-Cas systems (types I-C, I-F and III-A), two large prophage regions (Figure 3B) and a total of 85 STS in its I-C CRISPR arrays. In *B. producta* and *M. elsdenii,* the wealth of STS hit mostly in and around prophage regions, with some prophages remaining untargeted. After manual confirmation of the consensus repeat and array orientation of the STS, we observed that the oldest STS (located further from the leader in the CRISPR array) are those with protospacer in the prophage regions (Figure 3A and 3B), suggesting these were the initial hits and that additional spacers could have been acquired from locations in the prophage vicinity by primed CRISPR adaptation. Interestingly, as priming is enhanced by CRISPR interference (18,64,65), it is striking that no apparent DNA damage was incurred. For *M. elsdenii* we found that all STS protospacers are on the same strand with an orientation bias characteristic of primed adaptation (18). Primed adaptation would result in the acquisition of many spacers, explaining the high number of STS found in these genomes. It is interesting that STS in *M. elsdenii* were integrated in only two out of six CRISPR arrays, both close to the I-C *cas* genes (Figure 3B). It is also curious to note that no homologs of any known Acr (29) could be found in either genome using BLASTp homology searches with an e-value cutoff of 10^−5^.

Overall, these examples of extensive, tolerated self-targeting suggest that prophage integration was followed by primed adaptation, leading to the amplification of STS against the prophage and flanking genomic regions.

## DISCUSSION

Self-targeting CRISPR spacers (STS) in bacteria are not a rare phenomenon, as one fifth of bacteria with CRISPR systems carries STS. Interestingly, some types of CRISPR-Cas systems (i.e. types I-B and I-F) seem to be more prone to incorporation of STS into CRISPR arrays. As STS may lead to auto-immunity, here we questioned which mechanisms could drive STS acquisition and whether bacteria encode mechanisms to protect themselves. We observed a striking prevalence of prophages in STS-containing genomes when compared to STS-free genomes, suggesting that prophages could be the trigger of STS acquisition. Only about half of the STS targeted protospacers are located within the prophage regions, with the other half targeting the endogenous genome. Interestingly, STS hits in the endogenous genome are enriched in the proximity of prophages, showing a pattern consistent with primed adaptation from an initial protospacer present on the prophage. Also, in cases where bacteria carried multiple STS, the STS located the furthest from the leader sequence targeted the prophage, while subsequent STS targeted both prophage and endogenous regions. These results are consistent with a model where primed adaptation amplifies STS by acquisition of new spacers from both prophage and prophage-adjacent regions.

STS can lead to lethal auto-immunity, but we still found many STS-containing bacteria in the genome database, as well as many STS functional for direct interference (associated with a consensus PAM) capable of efficient targeting, especially in type I CRISPR-Cas systems. This suggests bacteria employ other mechanisms of auto-immunity evasion to survive. Interestingly, degradation of the CRISPR-Cas system itself does not seem to be the dominant evasion mechanism employed by bacteria to survive potential auto-immunity caused by STS, as we found at least 4 times more genomes with intact rather than degraded CRISPR-Cas systems. Genomes carrying type II and V CRISPR systems commonly have non-consensus PAM sequences of the STS protospacer which may help avoid auto-immunity. Whether this occurs by incorrect acquisition of the spacer (66,67), or mutation of the PAM when it is already integrated, is unknown. Although found at low frequency, Acrs were also present significantly (36-fold) more often in STS-containing genomes than STS-free genomes.

Based on our overall observations, we here suggest two scenarios for the appearance of STS in bacterial genomes. In the first scenario, bacteria may acquire a first spacer against a temperate phage, but despite this, the phage may still be able to integrate into the genome. In the second scenario, a prophage may already be integrated into the genome and the ‘accidental’ acquisition of an STS by the host may start targeting the prophage. Following this first STS, incomplete targeting may lead to further STS expansion by primed spacer acquisition, in type I and II systems (68,69), which will result in the incorporation of multiple new spacers targeting both the prophage and adjacent locations in the bacterial genome. This continuation of extensive priming, which is thought to require a level of CRISPR targeting, is without apparent genome damage or lethality. The process of acquiring STS creates an apparent standoff between CRISPR-Cas and targeted prophages that involves mechanisms of auto-immunity avoidance and anti-phage defense. As shown here, these interactions may involve Acrs that may contribute to creating tolerance to STS in general, and to STS-targeted prophages in particular. Thus, it is possible that the CRISPR system may be preventing prophage induction (70), and perhaps induce prophage clearance or genome deletions (71,72). When the protospacer region of the prophage in the bacterial genome is deleted, this may lead to interesting eco-evolutionary dynamics, as the presence of the former STS on the bacterial genome may now prevent reinfection of the immunized strain by the same or related phages. Similarly, if the CRISPR system prevents induction of the prophage by targeting it upon excision from the genome, the induction of the lytic cycle could be inhibited and the shift from lysogeny to a lytic state could be detected and acted upon. The balance between these processes remains subject to further experimentation and modelling.

It has been suggested that CRISPR-Cas systems could have some tolerance to mobile genetic elements to allow acquisition of potentially beneficial genetic information (73). Tolerance to prophages has been observed, but not to plasmids (73,74). Maintenance of a plasmid bearing beneficial traits in specific environmental contexts has been shown to lead to CRISPR loss (75,76), although probably resulting from the beneficial plasmid helping select for cells without CRISPR-Cas that could randomly appear in the population rather than the plasmid actively causing CRISPR-Cas loss. Tolerance may not be equal to all mobile genetic elements, such as mobile genetic elements that integrate the bacterial chromosome (e.g. prophages and transposons) as a consequence of the presence of Acrs or of selection for different modes of escape from self-targeting. Tolerance to integrated mobile genetic elements derived from auto-immunity escape may breach the barrier imposed by CRISPR-Cas systems and facilitate the diversification and evolution of bacterial genomes and the passive dissemination of phages in bacterial populations.

## AVAILABILITY

Data is available in the GitHub repository (https://github.com/hwalinga/self-targeting-spacers-scripts and https://github.com/hwalinga/self-targeting-spacers-notebooks).

## SUPPLEMENTARY DATA

Supplementary Data are available at NAR online.

