## Supplementary Figures for "Prophages are associated with extensive CRISPR-Cas auto-immunity"

### Content

**Supplementary Figure S1.** Distribution of self-targeting spacer length for each CRISPR-Cas subtype.

**Supplementary Figure S2.** *p*-value histogram of binomial test of the hypothesis that leader or tail STSs are more common than other STSs.

**Supplementary Figure S3.** STS position in CRISPR arrays of 10 or less spacers.

**Supplementary Figure S4.** Distribution of self-targeting spacer hits on the bacterial endogenous genome as a measure of distance to the nearest prophage as normalized by contig length.

**Supplementary Figure S5.** Distribution of the number of STS per genome of each CRISPR-Cas type.

**Supplementary Figure S6.** Prevalence of STS-containing genomes with *aca* genes as found by homology search to known *aca* genes.

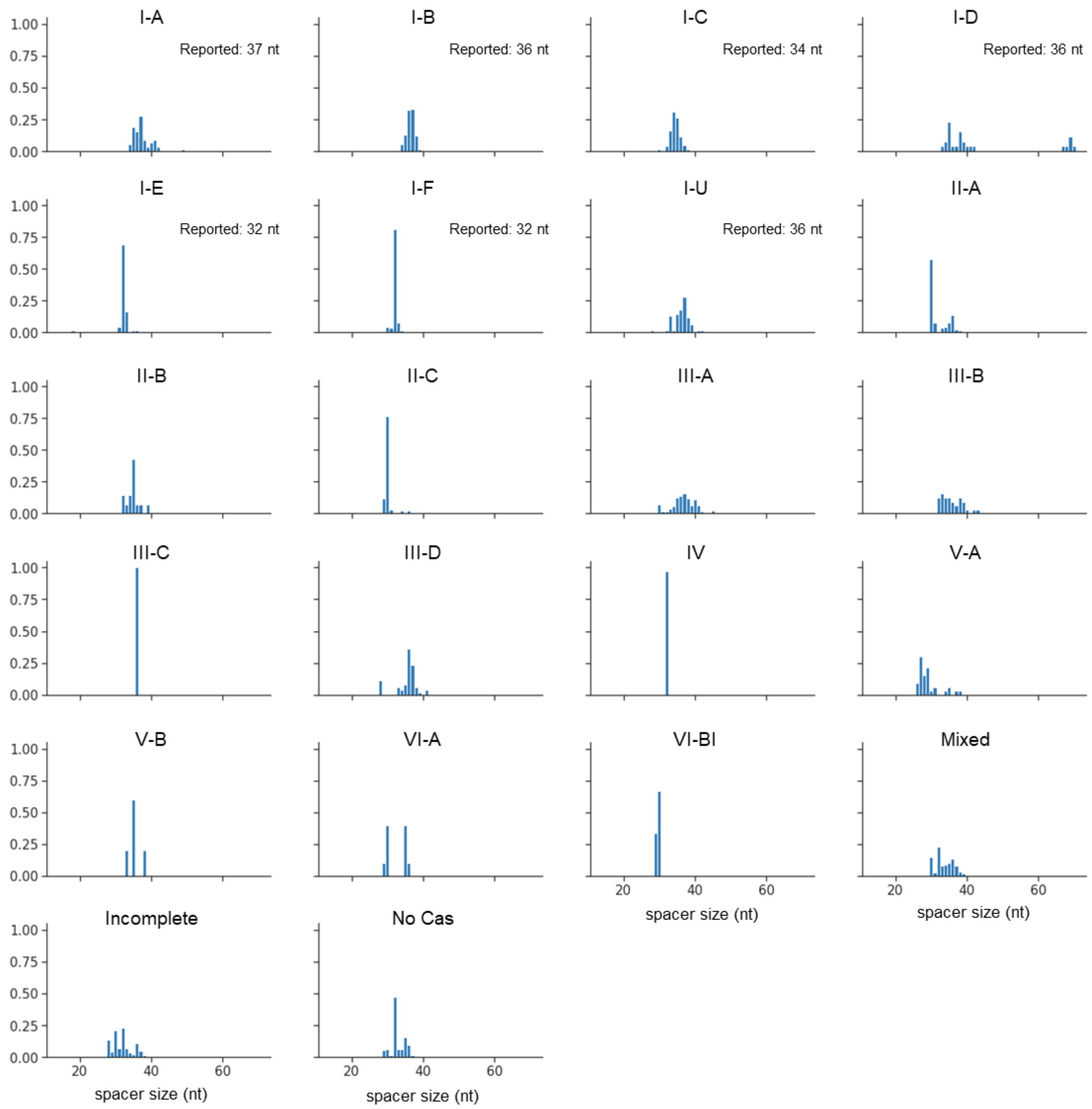


**Supplementary Figure S1**. Distribution of self-targeting spacer length for each CRISPR-Cas subtype. Reported preferred spacer length is indicated when known.

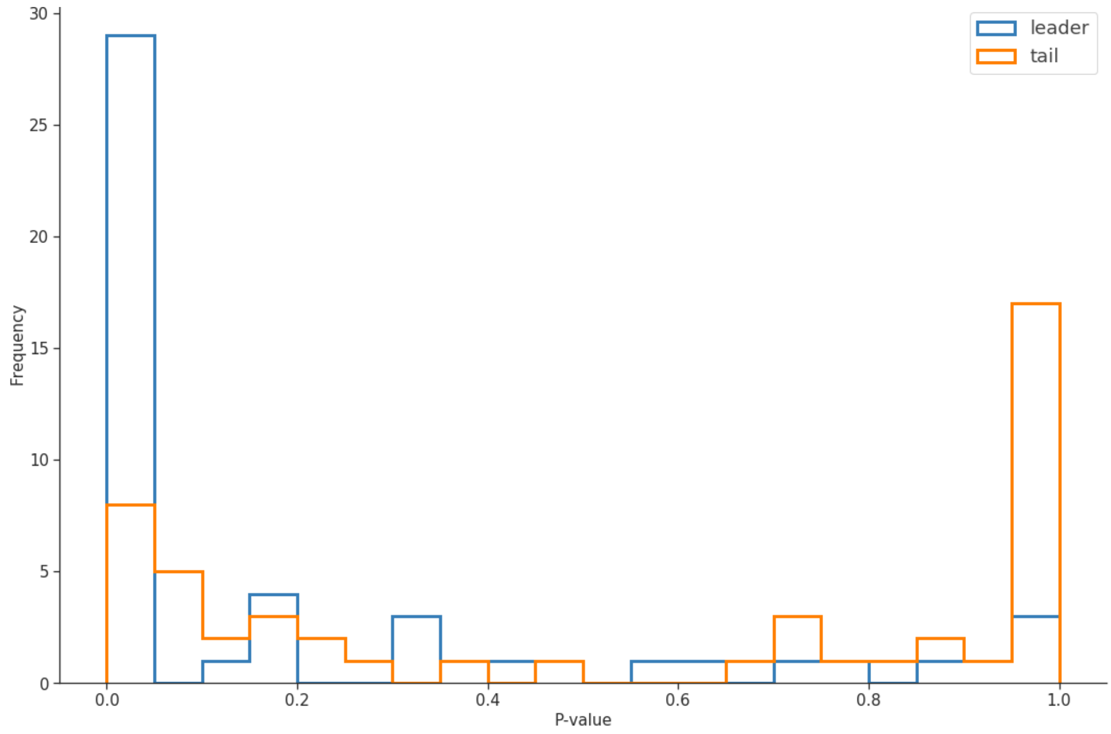


**Supplementary Figure S2.** *p*-value histogram of binomial test of the hypothesis that leader or tail self-targeting spacers (STSs) are more common than other STSs. Only STSs from CRISPR arrays smaller than 50 spacers were considered for this analysis. A larger bar in the $p < 0.05$ region indicates enrichment of STSs closer to the leader (blue). A larger bar in the $p = 1$ region indicates no enrichment of STSs closer to the tail (orange).

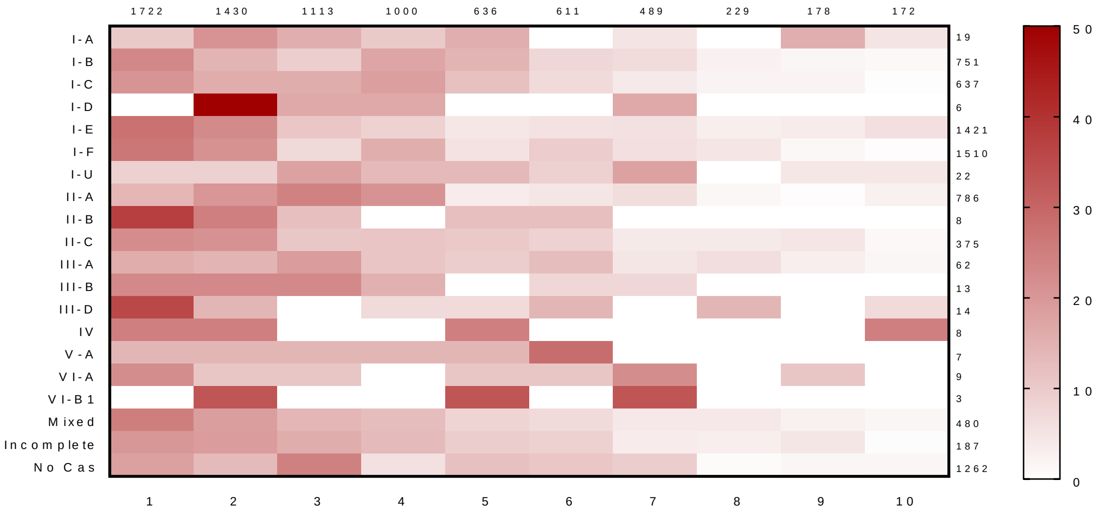


**Supplementary Figure S3.** STS position in CRISPR arrays of 10 or less spacers. Heatmap of STS position in the CRISPR array for each CRISPR-Cas subtype, using corrected orientation of the CRISPR arrays. Scale bar represents percentage of STS found per position in the CRISPR array. Total number of STS analyzed per CRISPR-Cas subtype is given for each row, while total number of STS per position in the array is given for each column.

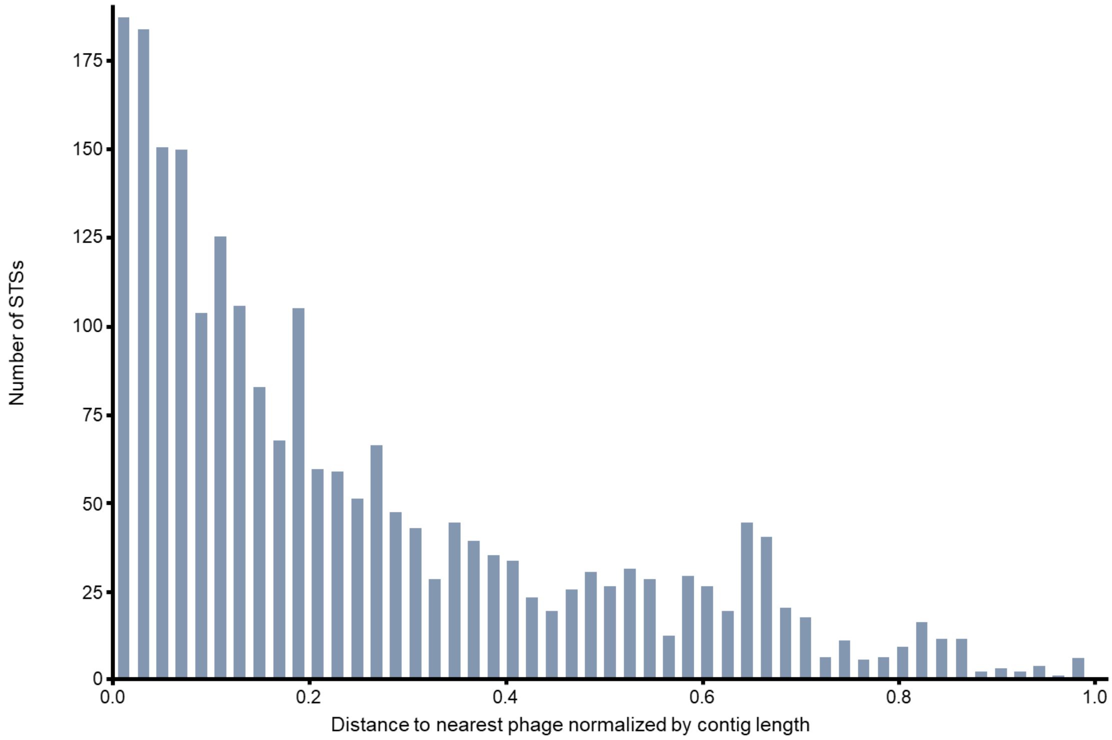


**Supplementary Figure S4.** Distribution of self-targeting spacer hits on the bacterial endogenous genome as a measure of distance to the nearest prophage, as normalized by contig length.

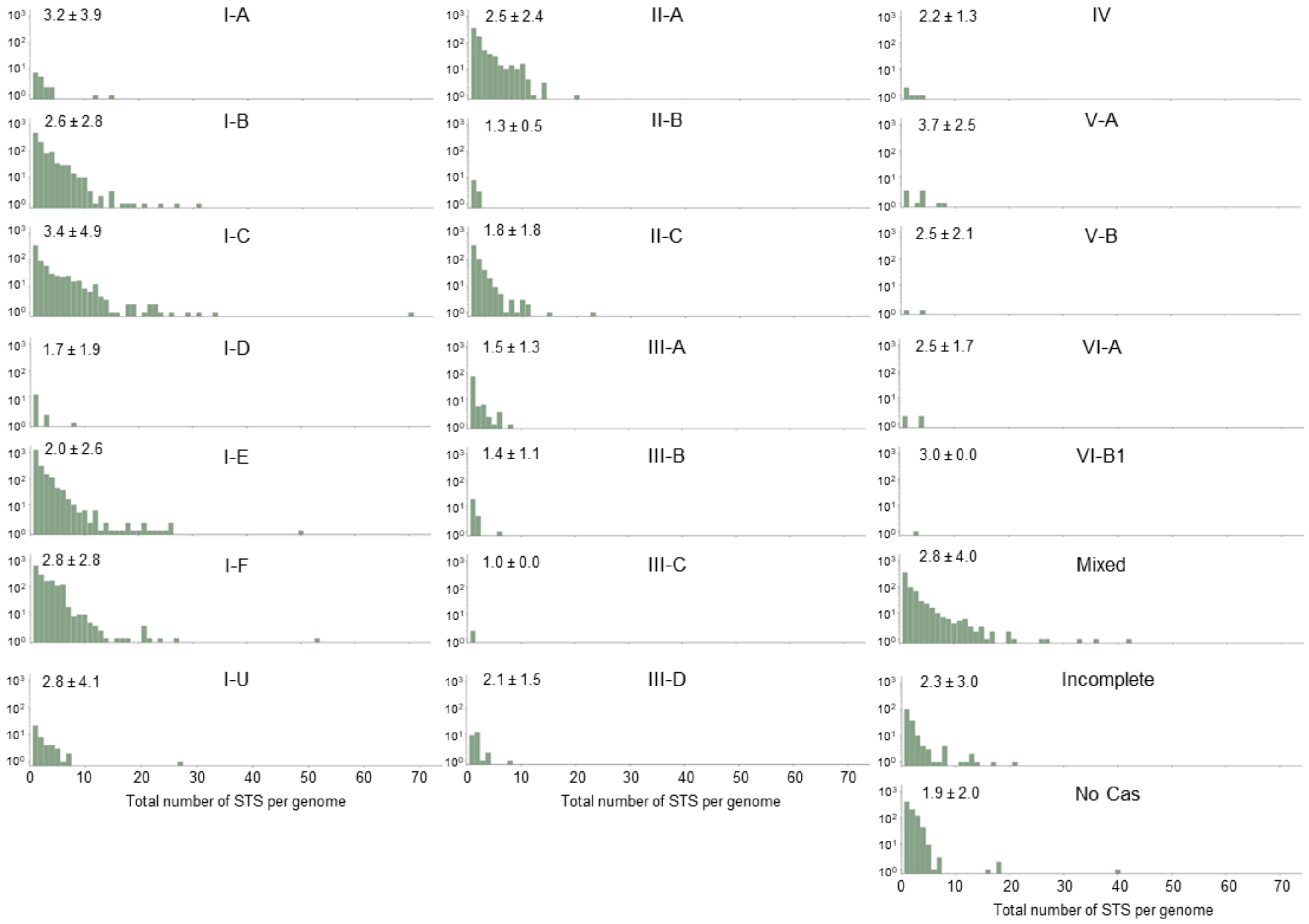


**Supplementary Figure S5.** Distribution of the number of unique STS per genome for each CRISPR-Cas type. Average number and standard deviation of STS per genome is indicated for each CRISPR-Cas type.

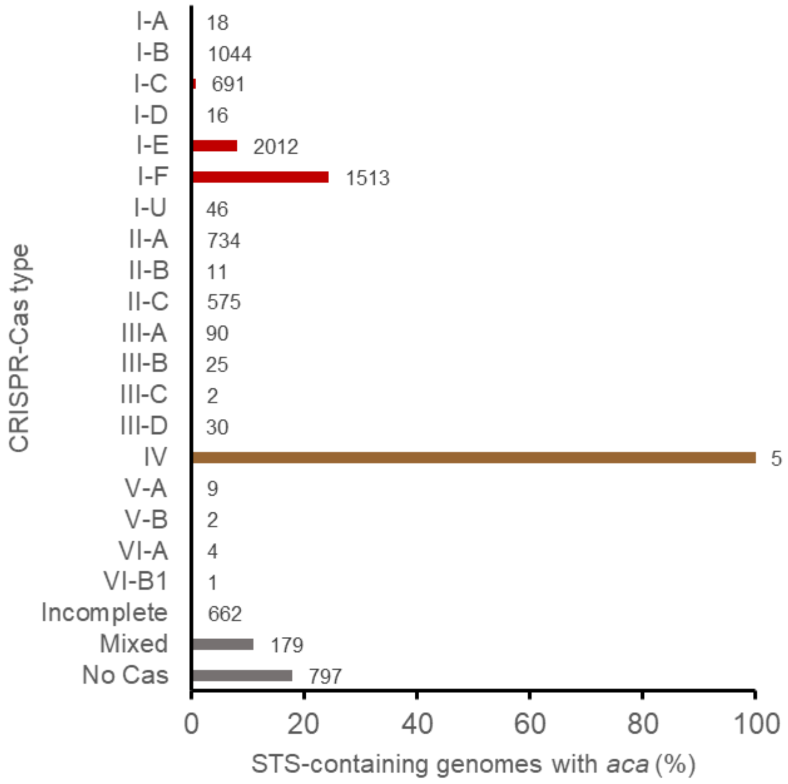


**Supplementary Figure S6.** Prevalence of STS-containing genomes with *aca* genes as found by homology search to known *aca* genes.
